# Viral Invasion Fitness Across a Continuum from Lysis to Latency

**DOI:** 10.1101/296897

**Authors:** Joshua S. Weitz, Guanlin Li, Hayriye Gulbudak, Michael H. Cortez, Rachel J. Whitaker

## Abstract

The prevailing paradigm in ecological studies of viruses and their microbial hosts is that the reproductive success of viruses depends on the proliferation of the “predator”, i.e., the virus particle. Yet, viruses are obligate intracellular parasites, and the virus genome – the actual unit of selection – can persist and proliferate from one cell generation to the next without lysis or the production of new virus particles. Here, we propose a theoretical framework to quantify the invasion fitness of viruses using an epidemiological cell-centric metric that focuses on the proliferation of viral genomes inside cells instead of virus particles outside cells. This cell-centric metric enables direct comparison of viral strategies characterized by obligate killing of hosts (e.g., via lysis), persistence of viral genomes inside hosts (e.g., via lysogeny), and strategies along a continuum between these extremes (e.g., via chronic infections). As a result, we can identify environmental drivers, life history traits, and key feedbacks that govern variation in viral propagation in nonlinear population models. For example, we identify threshold conditions given relatively low densities of susceptible cells and relatively high growth rates of infected cells in which lysogenic and other chronic strategies have higher potential viral reproduction than lytic strategies. Altogether, the theoretical framework helps unify the ongoing study of eco-evolutionary drivers of viral strategies in natural environments.

---

Viral infections begin with the physical interaction between a virus particle (the “virion”) and the host cell. Infection dynamics within the cell often culminate in lysis, i.e., the active disruption of the integrity of the cell surface, leading to the death of the host cell and the release of infectious virus particles [1, 2]. Virus-induced lysis can be a significant driver of microbial mortality at population scales[3–5]. As a result, studies of the ecological influence of viruses of microorganisms in natural environments have, for the most part, emphasized the impact of the lytic mode of infection. However, the spread of viruses through microbial populations need not involve the immediate lysis of the infected cell.

Indeed, many viruses have alternative strategies. Temperate phage – like phage *λ* – can integrate their genomes with that of their bacterial hosts, such that the integrated viral DNA, i.e., the prophage, is replicated along with the infected cell, i.e., the lysogen [6]. Chronic viruses, like the filamentous phage M13, infect cells and persist episomally [7, 8], whereby the genome is replicated and then packaged into particles which are released extracellularly without necessarily inducing cell washout [9, 10]. An analogous mode of chronic infection has been observed in archaeal virus-host systems [11]. These examples raise a critical question (see [12–14]): are temperate or chronic modes prevalent or rare in nature?

More than a decade ago, studies of marine, hydrothermal, and soil environments suggested that lysogeny could be more prevalent than assumed based on culture-based analysis of virus-microbe interactions [15–18]. This evidence has been augmented by recent studies identifying viral dark matter-including integrated and extrachromosomal viral sequences - in microbial genomes [19–21]. Yet, despite increasing evidence of the relevance of persistent infections *in situ* the ecological study of phage has not integrated a common metric to compare the context-dependent fitness of lytic, temperate, and other chronic viral strategies.

A landmark theoretical study provides a setting off point for investigating potential benefits of non-lytic strategies [22]. Using a combination of simulations and local stability analysis, this study proposed that temperate phage could persist over the long term if prophage integration directly enhanced host fitness or enhanced resistance to infections by other lytic phage (“superinfection immunity”). The same study predicted that oscillations in host population abundances could provide an ecological “niche” for temperate phage. In essence, if bacterial densities were too low to support the spread of lytic phage, then temperate phage already integrated into lysogens could persist until “conditions become favorable for the bacteria to proliferate” [22]. Yet this finding does not exclude the possibility that lytic strategies could out-compete temperate strategies – even if lysis at low densities leads to population collapse.

More recently, efforts to understand why viruses should be temperate have drawn upon the mathematical theory of bet hedging [23]. According to this application of bet hedging theory, the temperate strategy enables viruses to expand rapidly during stable periods for hosts (via lysis) and mitigate risks of population collapse, particularly during unfavorable periods for hosts (via lysogeny). Such estimates of long-term growth rates are limited in their applicability as they do not include explicit dynamics of infected cells nor subsequent virus-microbe feedback. Moreover, a focus on *long-term* estimates of growth does not address whether or not lysis is the advantageous strategy for a virus at a given *moment* in time. As noted by [23], ecological models that incorporate explicit mechanisms underlying virus-host interactions are required to understand the viability of viral strategies.

In this paper, we draw upon the foundations of mathematical epidemiology to quantify viral fitness measured in terms of the proliferation of *infected cells* instead of virus particles. In doing so, we propose to adapt the basic reproduction number, *𝓡*_0_, “arguably the most important quantity in infectious disease epidemiology” [24] to analyze broad classes of virus-microbe dynamics, including those characterized by lytic, latent, and chronic strategies. In particular, our study unifies previous proposed definitions for purely lytic phage (i.e., termed the “phage reproductive number” [25–27]) and application of the *𝓡*_0_ concept to temperate phage (where virion production requires cell lysis) [28, 29], while also extending the scope of applicability to chronic viral strategies (where virion production does not require cell lysis). Further, we show how using a unified metric to characterize viral invasion dynamics can help predict and explain a continuum of infection strategies observed in different environmental contexts.

## Results

### Cell-centric metric of viral fitness

We consider the spread of viruses through a microbial population. Foundational work in virus-microbe dynamics conceptualized viruses as “predators” and bacteria as their “prey” (sensu [30, 31]) – a paradigm that has become well-established with time (e.g. [32]). Rather than applying the conventions of predator-prey theory, we analyze models of virus-microbe dynamics in terms of the spread of an infectious disease (see Figure 1). The spread of an infectious disease can be quantified in terms of the basic reproduction number, *𝓡*_0_, i.e., when *𝓡*_0_ *>* 1 then a pathogen is expected to increase its relative abundance in a population [33, 34]. Here we propose the following definition of the basic reproductive number for generalized virus-microbe dynamics:

> 𝓡_0_: the average number of new infected cells produced by a single (typical) infected cell and its progeny virions in an otherwise susceptible population.

**FIG. 1.**
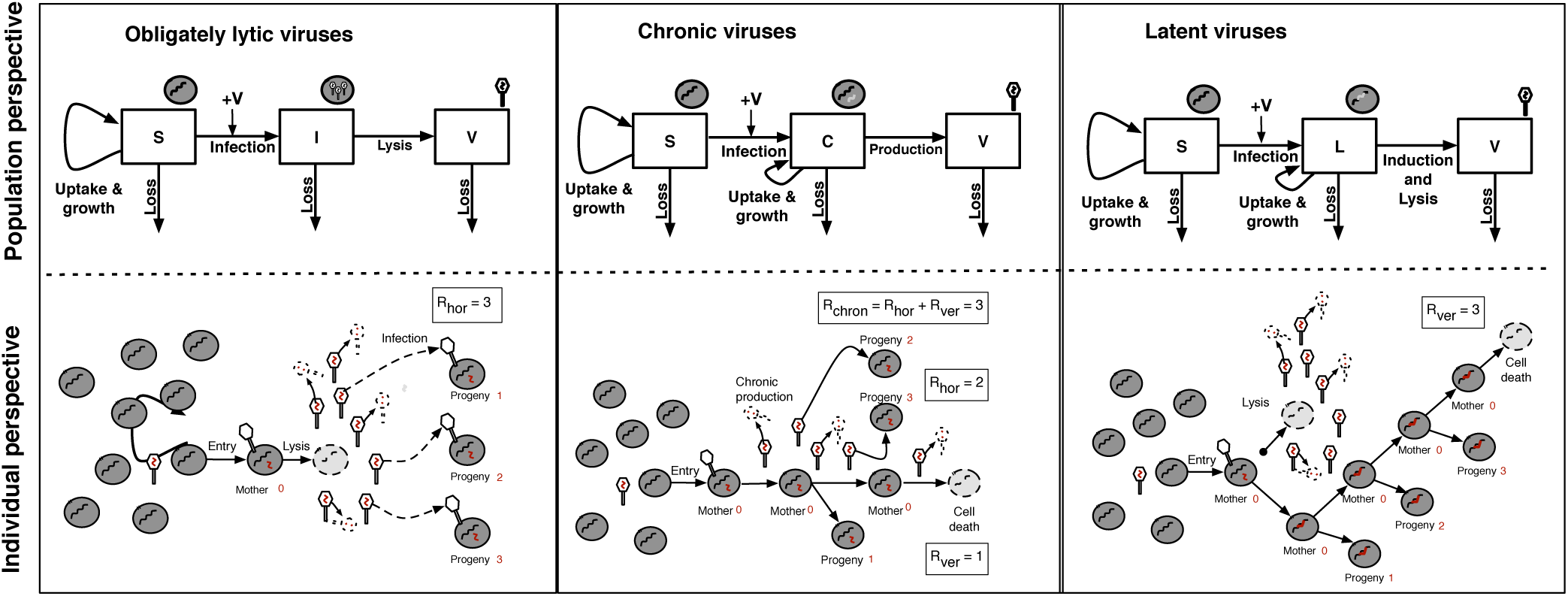
Schematic of obligately lytic, chronic, and latent strategies, give a population perspective (top) and individual perspective (bottom). (Top) The nonlinear dynamics for each model are presented in the Methods. (Bottom) The basic reproduction number accounts for complete cycles beginning with infected cells. In the obligately lytic case, only three virions of many infect cells, these are progeny, aka new mothers. In the chronic case, the mother cell divides once and two progeny virions infect new cells. In the latent case, the mother cell divides three times before it is removed. In all of these examples, the total reproduction number is the same, albeit with differing contributions from horizontal and vertical transmission routes.

This definition of *𝓡*_0_ counts viral reproduction in terms of infected cells, as in the study of eco-evolutionary dynamics of temperate phage [28, 29], rather than in terms of virus particles. In doing so, this definition builds upon insights from the virocell paradigm [35, 36], wherein the “real living [viral] organism” [35] is an infected cell actively reproducing new virions, i.e., the “virocell”. However, here we depart from a strict virocell definition, by accounting for transmission via virions and transmission via latent infections, e.g., where viral genomes are integrated into the genomes of their hosts.

As we will show, the cell-centric definition of viral fitness proposed here facilitates comparison of infections caused by “vertical” transmission (i.e., from mother to daughter cell), those caused by “horizontal” transmission (i.e. from an infected cell to another susceptible cell in the population), and those caused by a combination of both routes (e.g., as in chronic viruses). In doing so, *𝓡*_0_ quantifies fitness at the individual scale, i.e., beginning with the virus infection of a single microbe, and also represents a threshold condition for viral invasion at the population scale. Note that our definition of fitness does not account for feedback between viral population growth and the environment, an issue we return to in the Discussion.

### Obligately lytic viral strategies – a baseline for comparison

Dynamics of obligately lytic viruses and their microbial hosts can be represented via a set of nonlinear differential equations (see Figure 1 for this and other model schematics and Methods for equations). The spread of viruses in an otherwise susceptible population in Eq. (4) were analyzed using the Next-Generation Matrix (NGM) approach (see Materials and Methods). Via NGM, we find that obligately lytic viruses should spread when

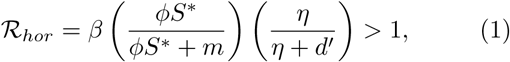

where *β* is the burst size, *φ* is the adsorption rate, *m* is the decay rate of virus particles, 1*/η* is the latent period, *d′* is the equi-librium density of susceptible cells. This threshold value represents the (exclusively) horizontal contributions to the basic reproduction number. This inequality can be understood in two ways (see Figure 1).

First, consider a single virion. Virions successfully adsorb to susceptible hosts at a rate *ϕS\**. In contrast, virions decay at a rate *m*. The factor *ϕS*/* (*ϕS\** + *m*) denotes the probability that a virion is adsorbed before it decays. Adsorption need not lead to lysis, instead given a lysis rate of *η* and a loss rate *d′* of infected cells, then only a fraction *η/*(*η* + *d′*) of infected cells will lyse and release virions before being washed out of the system. Finally, these two probabilities must be multiplied by the burst size *β*, i.e., the number of new virions released, to yield the average number of new infectious virions produced by a single virion in a susceptible host population. This product is equal to the basic reproduction number, *𝓡*_*hor*_ (what was previously termed the phage reproductive number [25–27]). When *𝓡*_*hor*_ exceeds 1 then a single virion produces, on average, more than one virion, of which each in turn produces, on average, more than one virion and so on. Figure 2 shows how viral proliferation varies with life history traits (in this case, the burst size) and the ecological context (in this case, the initial cell density). The critical value *𝓡*_0_ = 1 defines the threshold between regimes of viral extinction and proliferation.

**FIG. 2.**
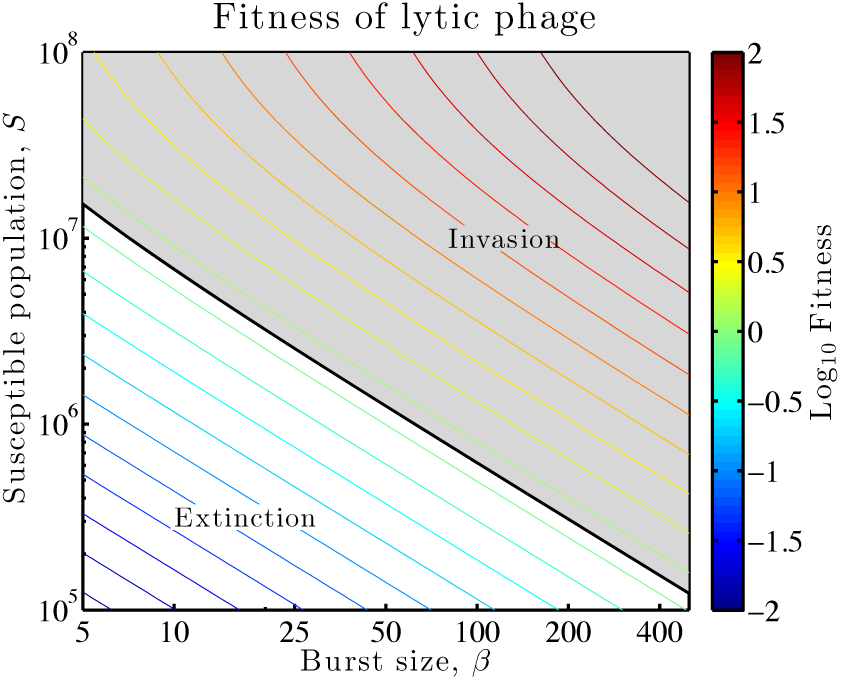
Virus reproduction as a function of burst size and susceptible cell density. The contours denote the log_10_ of *𝓡*_0_, as measured using Eq. (1), given variation in burst size, *β*, and susceptible cell density, *S*. Viruses invade when *𝓡*_hor_ > 1 or, equivalently when log_10_ *𝓡_hor_* > 0. Contours denote combinations of (*β, S\**) of equivalent *𝓡_hor_*. Additional parameters that affect viral reproduction are *ϕ* = 6.7 *×* 10^−10^ ml/hr and *m* = 1*/*24 hr^−1^.

Second, we can revisit this same calculation beginning with an assumption that there is a single infected cell in an otherwise susceptible population. In that event, the infected cell produces *β* virions a fraction *η/*(*η* + *d′*) of the time, of which only a fraction *ϕS*/* (*ϕS\** + *m*) are adsorbed before they decay. The product represents the number of newly infected cells produced by a single infected cell in an otherwise susceptible population. The product is the same, but in this alternative approach we have counted proliferation in terms of a viral life cycle that starts and ends inside cells. Although both interpretations – the virion-centric and the cell-centric – lead to equivalent estimates of *𝓡*_0_, we will use the cell-centric definition to unify subsequent comparisons across a spectrum of viral strategies.

## Latent viral strategies

In this section we consider the dynamics of latent viral strategies, such as temperate phage, in which proliferation may be either horizontal or vertical (but not both simultaneously). First, we focus on the case where virus genomes exclusively integrate with host cell genomes which can then be passed on to daughter cells. We use the cell-centric interpretation as before, and consider infection dynamics given a single lysogen in an otherwise susceptible population (see Eq. (5)). Via NGM analysis detailed in the Materials and Methods we find that lysogens proliferate when

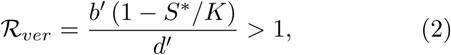

where *b*′ denotes the division rate of infected cells, *K* is the density of susceptible cells in the absence of viruses, and other parameters are equivalent to the obligately lytic model. Here the value of *R*_0_ is derived from *vertical* transmission of viral genomes among lysogens.

The basic reproduction number can be interpreted mechanistically. The term *b′*(1 − *S*/K*) represents the birth rate of lysogens, which decreases with increasing number of cells - whether susceptibles or lysogens. Given that *d′*is the death rate of lysogens, the term 1*/d′*denotes the average lifespan of an individual lysogen. Therefore, this reproduction number is equal to the average number of newly infectious cells produced in the lifetime of the original infection (see Figure 1). When *𝓡*_*ver*_ exceeds one, then a single lysogen will beget more than one lysogen, on average, and those lysogens will do the same, and so on.

As is evident, lysogens reproduce more frequently when they are subject to less competition with hosts, i.e., when *S\**is small relative to *K*. Given the value of *S\**, the basic reproduction number can be written as *𝓡_ver_* = (*b′/d′*) (*b/d*). Hence, if lysogens have more advantageous life history traits than do susceptible cells then viruses can spread exclusively via vertical transmission. This benefit of lysogeny applies in the immediate term and constitutes direct support for how a lysogen that benefits its host can also benefit the virus. However, if lysogeny comes with a cost (i.e., *b′/d′* lower than *b/d*), then vertical transmission alone will not be enough for *𝓡*_*ver*_ > 1. Note that *𝓡*_*ver*_ is a monotonically decreasing function of *S\**, such that increased abundances – all things being equal – diminishes the advantage for vertical transmission.

This analysis raises the question: does a strictly lytic or strictly lysogenic strategy have a higher basic reproduction number? Recall that the horizontal *𝓡*_0_ is an *increasing* function of susceptible cell density, i.e., when there are more hosts then the value of horizontal transmission increases. The value of *𝓡*_*hor*_ and *𝓡*_*ver*_ cross at a critical value, *S_c_* (see Figure 3). For *S > S_c_*, then horizontal transmission is favored and for *S < S_c_* then vertical transmission is favored.

**FIG. 3.**
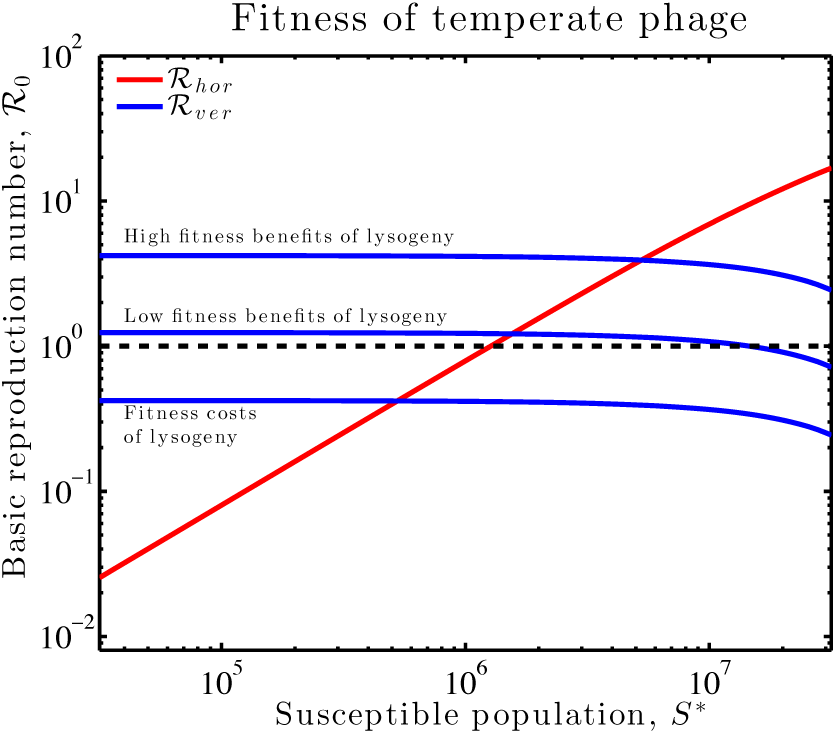
Basic reproduction number of temperate viruses as a function of susceptible cell density. The increasing (red) line denotes the horizontal *𝓡*_0_ if temperate phage infect then always lyse cells. The decreasing (blue) line denotes the vertical *𝓡*_0_ if temperate viruses always integrate with their hosts. Relevant parameters are *β* = 50, *ϕ* = 6.7 × 10^−10^ ml/hr, *K* = 7.5 × 10^7^ ml^−1^, and *b′* = 0.32, 0.54 and 1 hr^−1^ as well as *d′* = 0.75, 0.44, and 0.24 hr^−1^ for the three lysogeny curves from bottom to top respectively.

### Chronic viral strategies

Finally, we consider the dynamics of “chronic” virus strategies, or what have been termed “persister” or “producer” strains in other contexts (see Figure 1 and Methods). In a chronic infection both vertical and horizontal transmission can take place concurrently. Via a NGM analysis we find that a small number of chronically infected cells will spread in a population when

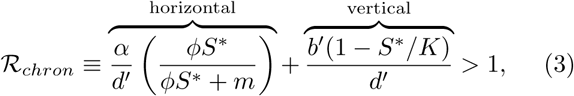

where *α* is the virus particle production rate and all other parameters are equivalent to those defined in the obligately lytic and and latent models. In the horizontal pathway, the chronic cell will remain viable for an average duration of 1*/d′*. In that time, the chronic cell will produce new virions at a rate *α*, of which only *ϕS*/* (*ϕS\** + *m*) will survive to enter another cell. Concurrently, the chronic cell will divide initially at a rate *b′*(*S*/K*), which when multiplied by the average cell duration of 1*/d′* yields the expected number of daughter cells, i.e., representing vertical transmission. A chronic virus will spread at the population scale due to the combination of transmission via horizontal and vertical components.

The spread of chronic viruses depends on both infected cell traits and virion-associated traits. As a consequence, it would suggest that chronic viruses should evolve to improve the sum of horizontal and vertical reproduction. Trade-offs likely constrain the evolution of virion release rates and cell duration. For example, increasing the virion production rate, *α*, may improve horizontal reproduction, but if doing so increases cell washout, *d′*, then the overall change in *𝓡*_*chron*_ may be negative. As a result, it is possible that chronic viruses could have the largest reproduction number in an intermediate density regime (see example in Figure 4). Understanding the pleiotropic effects of changes to chronic virus genotypes may provide one route to characterizing the evolution of viral strategies in which both horizontal and vertical transmission rates operate concurrently [37].

**FIG. 4.**
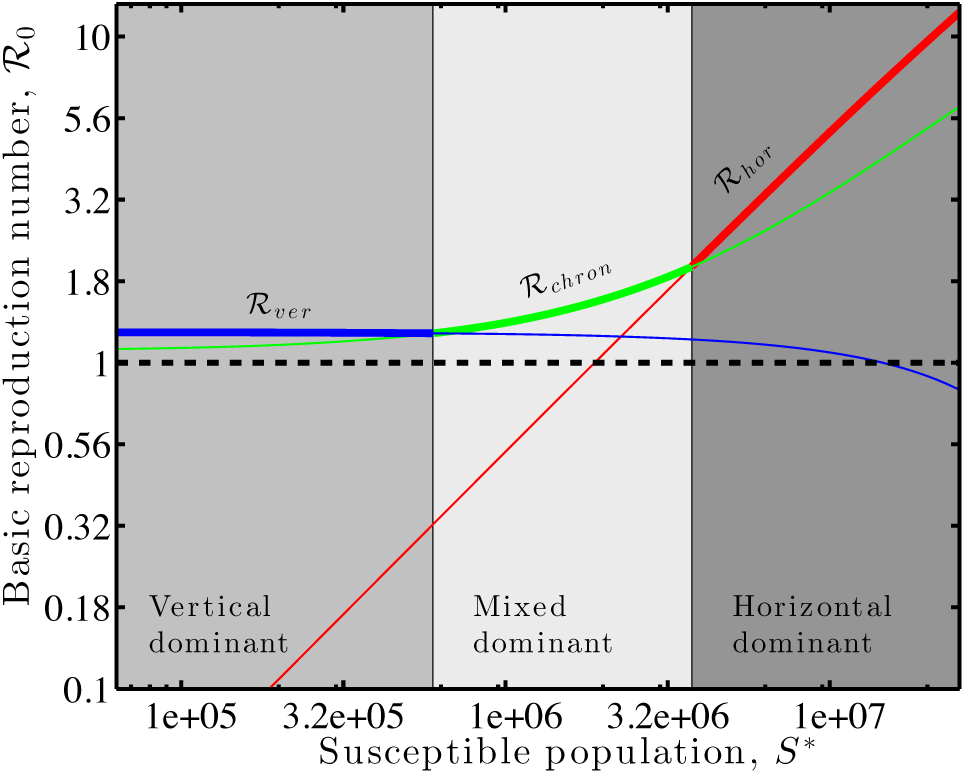
Viral strategies with the highest *𝓡*_0_ vary with susceptible host density, including exclusively vertical (bold blue, left), mixed (bold green, middle), and horizontal (bold red, right) modes of transmission. Relevant parameters are (i) for obligately lytic viruses (red): *β* = 100, *ϕ* = 6.7 × 10^−10^ ml/hr, and *m* = 0.13 hr^−1^; (ii) for chronic viruses (green): *b′* = 0.68 hr^−1^, *d′* = 0.63 hr^−1^, *α* = 20 hr^−1^, *ϕ* = 3.4 × 10^−10^ ml/hr, and *m* = 0.04 hr^−1^; (iii) for temperate viruses, given vertical transmission (blue) *b′* = 0.54 hr^−1^, *d′*= 0.44 hr^−1^, where *K* = 7.5 × 10^7^ ml^−1^ in all three scenarios given variation in *S\**.

## Discussion

We have proposed a unified theoretical framework to measure the spread of viruses within microbial populations when utilizing strategies spanning a continuum from lysis to latent to chronic. By defining viral reproduction in terms of infected cells, we are able to directly compare the spread of obligately lytic viruses, latent viruses, and chronic viruses in the context of nonlinear population models (see Figure 1). The invasibility of a newly introduced virus is measured in terms of the basic reproduction number, specifically adapted to the life cycle of viral infections of microbial hosts.

At its core, the theoretical framework re-envisions life history theory for viruses that infect microorganisms. In our calculations, a focal virus genome inside a cell can be thought of as a “mother virus”. These mother viruses may lyse cells and produce “juvenile” offspring, i.e., virus particles. When a virion successfully infects a susceptible host this new infection becomes, once again, a mother virus. This is an example of horizontal transmission. For latent and chronic viruses, the viral genome inside an infected cell may be passed on to both cells upon division. This division is equivalent to direct reproduction of a mother virus, bypassing the juvenile state. This is an example of vertical transmission. Combinations of these two scenarios emerge in applying next-generation matrix theory for calculating the basic reproduction number of viral strategies (see Figures 2-4 and the Materials and Methods).

The critical invasion fitness of a virus strategy, as calculated in terms of *R*_0_, depends on life history traits as well as susceptible cell density. Obligately lytic viruses have increasing values of *R*_0_ in populations with larger numbers of susceptible hosts. This trend is consistent with experimental findings that fitness of virulent phage *λ*cI857 declines with decreasing susceptible cell density [28]. More broadly, we contend that the link between strategy and susceptible host density will inform ongoing debates on conditions that favor lysogeny and other persistent infections in marine systems; debates that have focused on the ratio of free virus particles and total microbial densities [12–14, 38]. Our analysis makes it evident that evaluating the benefits of latent or chronic strategies also requires consideration of the intracellular infection status of hosts. For example, in our models, strictly (or partially) vertically transmitted viruses may be favored when new susceptible hosts are scarce and when infections benefit host competitive fitness, e.g., through growth or survival. This may help to explain the inverse relationship between inducible lysogen fraction and total microbial abundances in marine environments [39].

The present approach adapts an epidemiological framework beyond temperate phage (as analyzed by [28, 29]) to include obligately lytic and chronic strategies. In doing so, we have focused our analysis on short-term invasion scenarios. Comprehensive understanding of viral strategies requires analysis of both short- and long-term dynamics [40]. This is particularly relevant given that evolutionary dynamics need not lead to the maximization of *𝓡*_0_ (reviewed in [41]). Indeed, long-term fates are influenced by the Malthusian (i.e., exponential) growth rate of viruses which we denote as *r*. The basic reproduction number *𝓡*_0_ and *r* are related, but they are not equivalent [42] – *𝓡*_0_ measures the speed of viral proliferation in generations (i.e., at the individual level) whereas *r* measures the speed of viral proliferation in time (i.e., at the population level) [43].

The threshold condition *𝓡*_0_ *>* 1 indicates whether the population growth rate *r* is positive, but does not predict changes in fitness given viral-host feedback. For example, virus proliferation depletes susceptible hosts, thereby decreasing the ‘effective’ viral fitness of the obligately lytic pathway – an outcome concordant with prior findings from mathematical models and eco-evolutionary experiments involving temperate phage *λ* and *E. coli* [28]. In the work of [28], obligately virulent phage increased in relative number in environments initially dominated by susceptible cells, whereas temperate phage strains exhibited higher population level growth when suseceptible cells were subsequently depleted via lysis. Extrapolating to chronic strategies, we expect that viral production may shift from horizontal to vertical as viruses proliferate through a microbial population.

Systematic analysis of the evolution of viral traits spanning lysis, latent, and chronic strategies in an ecological context is likely to draw upon a substantial body of work on the evolution of virulence (e.g., [44–51]). Priority areas include the evolution of traits when viral particles can persist for long periods in the environment (similar to epidemiological models of the ‘curse of the pharaoh’ [52, 53]) and the evolution of transmission mode itself [54]. In moving forward, one immediate opportunity is to assess how viruses of microbes evolve virulence levels, or even strategy types, when co-infecting the same microbial population. For example, analysis via the cell-centric approach implies that lytic viruses may reduce niche competition between cells, increasing the benefits of vertical transmission, and enable invasion by latent or chronic viruses [55]. This finding is consistent with repeated evidence of coinfection in microbial genomes between ssDNA filamentous phage (*Inoviridae* that have a chronic lifestyle) and dsDNA phage (*Caudovirales*, that can transmit via lysis) [19]. In addition, interactions of multiple viruses within the same host cell could lead to emergent new feedback strategies. For example, temperate phage can exhibit plastic strategies in which infection outcome depends on the multiplicity of infection [29, 56–60]. However, virus-virus interactions may also extend beyond the cell, e.g., some SPbeta viruses modify the state of bacterial cells through the release of small molecules, thereby shifting decisions between lysis and lysogeny during proliferation [61, 62].

Altogether, the theory presented here provides an additional imperative to develop new measurement approaches to assess the entangled fates of viruses and cells. Measurements of the fitness of viruses with latent and chronic strategies should prioritize estimates of the life history traits of infected cells. Screening for viral genomes and their expression inside cells – whether integrated or persisting episomally – may reveal benefits of viral strategies that have thus far remained hidden when utilizing lysis-based assays or virion counts. By combining measurements and theory, we hope that the present framework provides new opportunities to explore how viruses transform populations, communities, and ecosystems.

## Materials and Methods

### Parameters

The three model variants we analyze include a set of common parameters as well as parameters unique to particular models. The parameters include *b* and *b′*(maximal cellular growth rates, hr^−1^), *K* (carrying capacity, ml^−1^), *ϕ* (adsorption rate, ml/hr), *d* and *d′*(cellular death rates, hr^−1^), *β* (burst size), *m* (virion decay rate hr^−1^), *η* (lysis rate, hr^−1^), *p* (scaling factor for lysis, 0 *< p <* 1), *q* (scaling factor for latency, 0 *< q <* 1), and *α* (virion production/budding rate, hr^−1^). Additional context on viral life history traits, including constraints and estimation methods, is described in [63].

### Nonlinear dynamics, lytic strategies

The coupled system of nonlinear ordinary differential equations (ODEs) For the lytic system are:

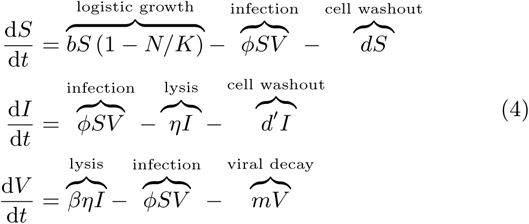

where *S*, *I*, and *V* denote the densities of susceptible cells, infected cells, and virus particles, respectively (see [63, 64]).

### Nonlinear dynamics, latent strategies

The nonlinear ODEs for the latent system are:

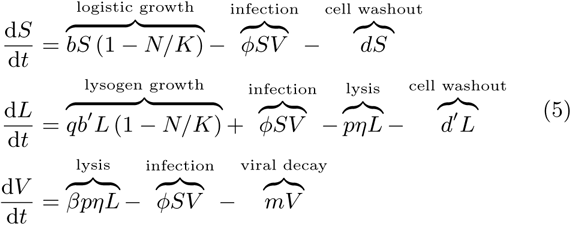

where *S*, *L*, and *V* denote densities of susceptible cells, lysogens (latent cells), and virus particles, respectively, and the total number of cells is *N* = *S* + *L*. Here, the relative rate of lysogenic growth and cellular lysis is controlled by the scaling factors *q* and *p*. When *q* = 1 and *p* = 0 then all infections are strictly latent and only lead to lysogenic growth. In contrast, when *q* = 0 and *p* = 1 then all infections are strictly lytic and only lead to cellular lysis. This is a variant of a nutrient-explicit formulation considered as part of an analysis of the tradeoffs underlying lysis and lysogeny for marine viruses [65]. Of note, this model does not include the absorption of virus particles to latent cells nor the degradation of prophage (i.e., “curing”).

### Nonlinear dynamics, chronic strategies

The nonlinear ODEs for the chronic strategies are:

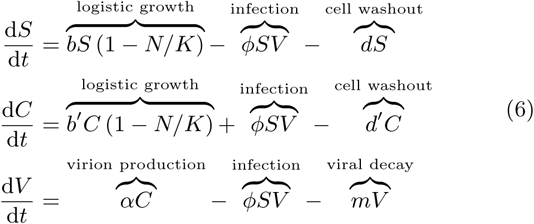

where *S*, *C*, and *V* denote densities of susceptible cells, chronically infected cells, and virus particles, respectively, and the total number of cells is *N* = *S* + *C*. Although the nonlinear population model of Eq. (6) be remapped to the latency model (Eq. (5)), this system of equations represents distinct mechanistic processes, including establishment of a chronically infected cell and release of virions from chronically infected cells without lysis at a per-capita rate *α*. Of note, this model assumes that infected mother and daughter cells both retain a copy of the viral genome. Partial fidelity of the vertical viral transmission process could be represented via a birth-dependent transfer of population from the *C* to the *S* states.

### Next-generation matrix (NGM)

We use the next-generation matrix (NGM) approach to calculate *𝓡*_0_ in math-ematical models of interactions between cells and viruses. We follow the convention of Dieckmann and colleagues in analyzing the subset of the epidemiological model including infected subclasses [34]. In the case of viruses of microbes, we denote those infected subclasses to include any population type that has an infectious viral genome, i.e., both infected cells and virus particles.

### NGM - Obligately lytic interactions

We linearize the dynamics of Eq.(4) around the virus-free equilibrium, (*S*\*,0,0) where *S*\* = [*K*(1 − *d/b*), and focus on the infected subsystem of *X*(*t*) = [*I*(*t*) *V* (*t*)]^┬^. The linearized infected subsystem dynamics can be written as 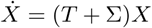 where

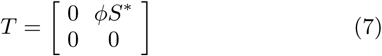

denote transmissions events (i.e., corresponding to epidemiological births) and

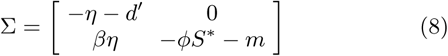

denote transmissions events (i.e., corresponding to changes in the state of viral genomes, including loss of infections). Via the NGM theory, the basic reproduction number *𝓡*_0_ corre-sponds to the largest eigenvalue of the matrix −*T*∑^−1^. The *i, j* matrix elements of Σ^−1^ correspond to the expected duration in state i of a viral genome that begins in state j. For this model,

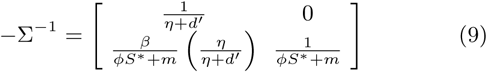

As a consequence, the basic reproduction number is:

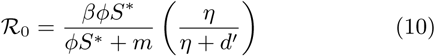

### NGM - Latent strategies

The linearized infected subsystem dynamics of Eq.(5) where *X*(*t*) = [*L*(*t*) *V* (*t*)]^┬^ can be written as 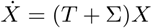 where

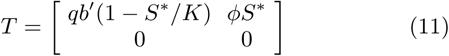

denote transmission events and

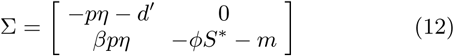

denote transition events. For this model,

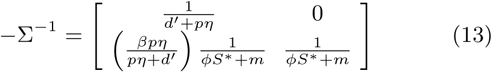

As a consequence, the basic reproduction number is:

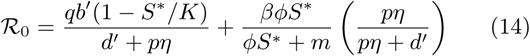

### NGM - Latent strategies

The linearized infected subsystem dynamics of Eq.(5) where *X*(*t*) = [*I*(*t*) *V* (*t*)]^┬^ can be written as 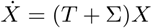 where

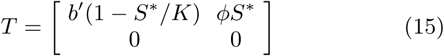

denote transmission events and

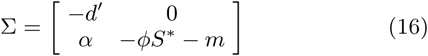

denote transition events. For this model,

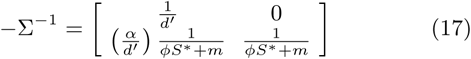

As a consequence, the basic reproduction number is:

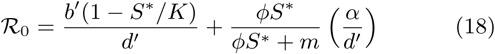

## Data availability statement

Simulation code is written in MATLAB and available at https://github.com/WeitzGroup/virfitness.

## Correspondence

Requests for materials and correspondence should be directed to JSW.

## Acknowledgments

JSW thanks J. Lennon, M. O’Malley, and the participants of a 2017 CIFAR-GBMF workshop on ‘A Continuum of Persistence’ for discussions critical to the development of ideas in this paper. The authors thank S. Brown, J. Dushoff, S. Elena, B. Levin, M. Sullivan, and S. Wilhelm for comments and feedback on the manuscript. This work was supported by a grant from the Simons Foundation (SCOPE Award ID 329108, J.S.W.), an Allen Distinguished Investigator Grant (to R.J.W.), and a NSF Dimensions of Biodiversity grant 1342876 (to J.S.W. and R.J.W.).

## Contributions

JSW and RJW designed the study. JSW, GL, HG, and MHC contributed analytic methods. JSW performed computational analyses. JSW, MHC, and RJW wrote the paper.

